# Master regulators of signaling pathways coordinate key processes of embryonic development in breast cancer

**DOI:** 10.1101/425777

**Authors:** Diana Tapia-Carrillo, Hugo Tovar, Tadeo E. Velazquez-Caldelas, Enrique Hernandez-Lemus

## Abstract

Signal transduction pathways allow cells to respond to environmental cues and can induce intracellular changes. In some contexts, like embryonic development, signal transduction plays crucial roles in cell fate determination and differentiation, while in developed organisms some of this processes contribute in the maintenance of the structural integrity of tissues.

Tumor cells are recognized as having deregulated signaling which leads to a series of abnormal behaviors known as the hallmarks of cancer. Although gene regulation is often viewed as the last step in signal transduction, transcriptional regulation of the components of a pathway may impact in the long term deregulation observed in tumors. The study of gene regulatory networks centered around genes of the signal transduction pathways allows the identification of transcriptional regulators with the greatest influence over the signal transduction gene signature, also denominated Master Regulators.

In this work we identify, the master regulators that regulate the expression of genes of 25 relevant pathways grouped in KEGG within the category of signal transduction in a breast cancer dataset. For this purpose we implemented a modified MARINa algorithm that identifies, from a network of regulons, those that possess more differentially expressed genes related to the process to be studied. We identified CLOCK, TSHZ2, HOXA2, MEIS2, HOXA3, HAND2, HOXA5, TBX18, PEG3 and GLI2 as the top 10 master regulators of signaling pathways in breast cancer. Nine of them are recognized for taking part in embryonic development associated processes.

Individual enrichment GO biological function for each TMR regulons showed to be significantly enriched in embryonic development related processes. Hedgehog signaling pathway was shown as enriched and also highly deregulated. The genes of the HOXA family are shared among most of the TMRs. Overall, this suggests the importance of the aberrant reprogramming of mechanisms present during embryonic development, being coopted in favor of tumor development.

## Introduction

Breast cancers are illnesses that originate from healthy cells that are somehow reprogrammed to acquire unlimited proliferation and self-renewal capacity. Eventually these transformed cells are able to migrate and invade other tissues in the body [1]. In this context, cancerous cells do not create brand new cell signaling pathways, but through a variety of mechanisms, existing pathways are aberrantly activated [2]. On them it is notable that many pathways associated to tumor development also play important roles during embryonic development. Pathways such as WNT, Hedgehog or VEGF are essential for differentiation, migration and pattern formation. Abnormal expression of components of these pathways have been reported in tumors [3] [4] [5].

Theories exist regarding the origin of cancerous cells and its relationship to embryonic development pathways and on how these contribute in dedifferentiation and later specialization on tumor development. Nevertheless, all of them agree on the presence of deregulation of signal transduction pathways (STP) controlling these processes [6] [7]. Accounting for gene expression as one possible mechanism in the modulation of signal transduction pathways, the determination of transcription regulators may help us understand this phenomenon.

A significant level of regulation of signaling is achieved through the action of transcription factors (**TFs**) that modulate the transcription of groups of genes encoding proteins that participate in these pathways [8–10]. Given their capacity to modulate cellular pathways, TFs are of great interest in the study of complex diseases [11] [12] [13]. Moreover, it has been recognized that some TFs exert a decisive influence in the transition between phenotypes. These TFs, called Transcriptional Master Regulators (TMRs) [14] are expressed at the early onset of the development of a particular phenotype, consequently regulating multiple target genes either directly or indirectly by means of transcription cascades resulting in significant gene expression changes and hence phenotype variation.

Given the fact that multiple signal transduction factors are simultaneously deregulated in the cancerous phenotype, an integrative approach is valuable in order to understand the biology underlying this disease. MARINa (Master Regulator Inference Algorithm) can infer TFs with greater influence in the transition between healthy and diseased phenotype in genetic regulation networks of the breast cancer phenotype [14,15]. In this work, we used a modified version of this algorithm to find the most important transcription factors focused in the regulation of KEGG’s 25 signal transduction pathways in breast cancer. We also identified a TMR subset that regulates genes belonging to specific signal transduction pathways in breast cancer.

## Materials and methods

### Obtaining and preprocessing data

A Gene Expression matrix was obtained from Espinal-Enriquez *et al.* [16]. Corresponding to The Cancer Genome Atlas (TCGA) level 3 available data of the Illumina HiSeq RNA-Seq platform, and consisting of 881 samples of which 780 correspond to breast cancer tissue and 101 to adjacent healthy mammary tissue. Quality control and batch effect removal were performed with *NOISeq* [17] and *EDASeq* [18] R libraries respectively [16].

### The Master Regulator Inference Algorithm

TMRs were inferred using the Master Regulator Inference Algorithm (MARINa) [15]. MARINa identifies TMRs through an enrichment of TF regulons (a TF with its targets) with differentially expressed genes between two phenotypes (breast cancer vs adjacent healthy mammary tissue). TMR inference with MARINa requires as input a network of regulons, a gene expression molecular signature and a null model [15] (Fig. 1). The construction of these elements is described below.

**Fig 1.**
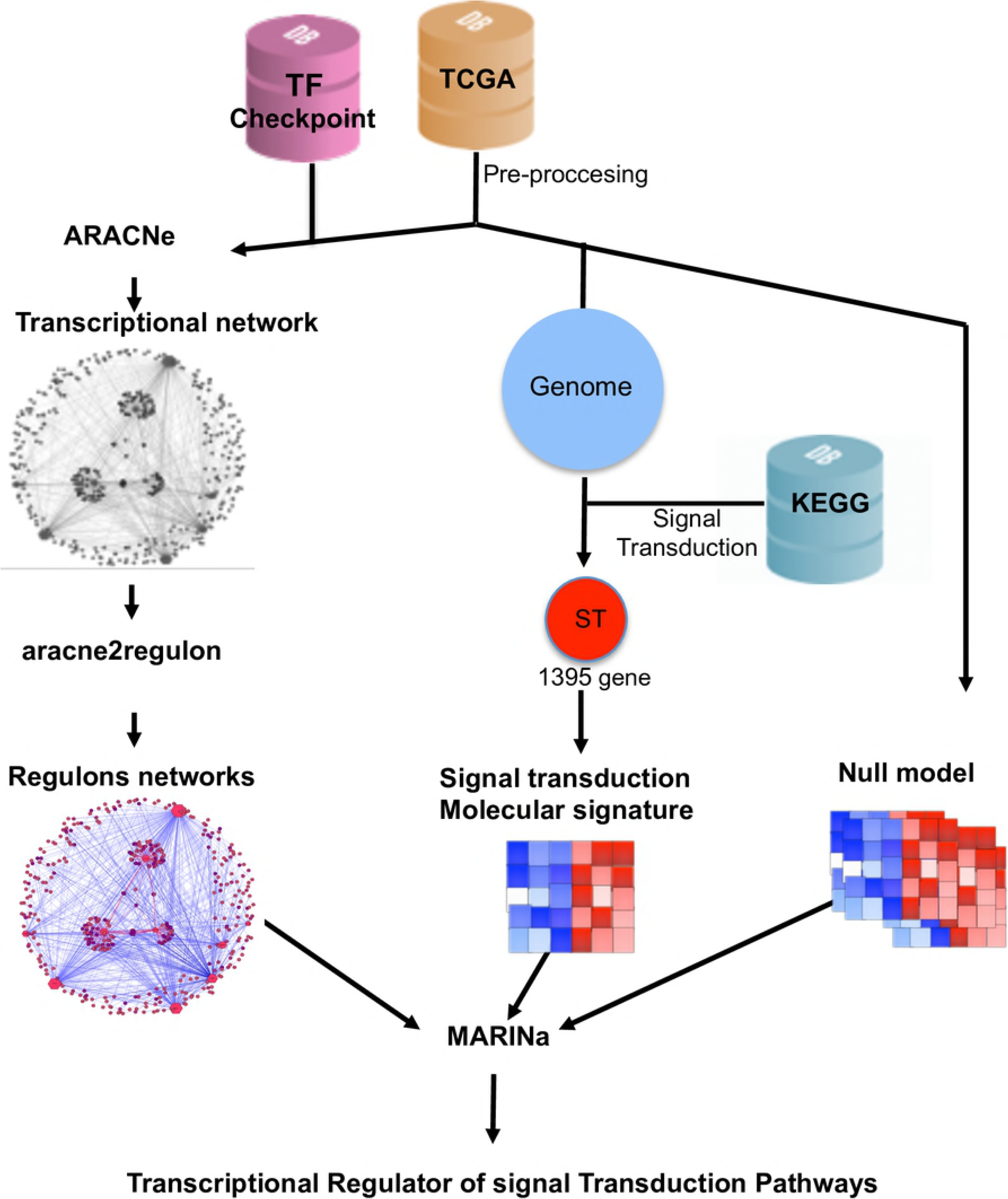
Pipeline. RNAseq data from TCGA’s 780 invasive mammary carcinomas and 101 adjacent tissue samples was processed to obtain an expression matrix (orange cylinder). The expression matrix and a list of transcription factors from the TFCheckpoint database (pink cylinder) served as input to infer a transcriptional regulatory network with ARACNe. A regulon network was obtained associating the expression level of the targets of all transcription factors using the *aracne2regulon* function from *viper* (left side). For the generation of the molecular signature, we considered genes in the expression matrix in KEGG’s ”signal transduction” category (blue cylinder). Finally, a null model was generated by permuting sample labels and recalculating the molecular signature (right). These three elements are the input to MARINa for the inference of the transcriptional master regulators (TMR) of the signal transduction pathways.

#### Generation of the regulons network

The network of regulons is a directed network (TF→Target) of all the transcription factors and their targets. To obtain it, we used the expression matrix and the mutual information based transcriptional regulation network built with ARACNe [19]. For this network, we considered transcription factors in the TFCheckpoint curated database [20] that possessed experimental evidence for TF activity. 771 of these TFs were found within the expression matrix S1 File.

This network contains the relationships between transcription factors and the rest of the genes, measured by the mutual information (MI) function [19,21]. For this network interactions were kept if its *p* value was below 0.005. Given that mutual information can detect both indirect and direct relationships, ARACNe limits the number of indirect interactions applying the Data Processing Inequality theorem (DPI), which considers that, in a triangle of interactions, the weakest one has a greater probability of being indirect if its difference is large with respect to the other two [22]. We applied a DPI value of 0.2 as recommended in Margolin *et al.* 2006 [19], which means that the weakest interactions of the triangles in the network were eliminated without introducing an excessive number of false positives.

The type of association (activation or repression) of the transcription factors is determined from Spearman correlation of the TF with the levels of expression of all its targets [15] this calculation was performed by the *aracne2regulon* function in the *viper* [23] R package.

#### Molecular signature generation of signal transduction pathways

In the standard MARINa workflow, the molecular signature is built by comparing the expression level distributions for all genes between two conditions (*e.g.*, healthy and diseased). For this work we built a molecular signature using only those genes annotated within the *signal transduction pathways* category in the Kyoto Encyclopedia of Genes and Genomes (KEGG) database [24]. For human, this category comprises 25 pathways. The total number of genes present in this subset is 1,700 of which 1,395 coincided with our expression matrix S2 File. The purpose of this filtering is to focus our search on those transcription factors that regulate the activity of these STPs in breast cancer. The molecular signature was built by applying a *t* test for each gene of the expression matrix, between tumors and adjacent healthy mammary tissue. The results of this test were *Z*-score normalized to allow comparability [15].

#### Null model generation

To estimate the probability that a Gene Enrichment Score depends on the biological context and thus is not merely random, a null model was generated by random permutation of samples between cases and controls and recalculation of differential expression [15].

#### Inferring the Master Regulators of signal transduction pathways

With the molecular signature, the regulon network and the null model, MARINa estimated the top regulons that enrich the most differentially expressed genes in the molecular signature through a Gene Set Enrichment Analysis [25]. An additional constraint was to consider only TFs with 20 or more targets in the molecular signature [15]. A *p*-value for each regulon was estimated by evaluating the Enrichment Score (ES) with reference to the distribution of scores of the null model [15]. For TMR inference we used Bioconductor’s *viper* package [23].

### Regulon enrichment of KEGG pathways

An over-represented pathway is defined as one for which we found significantly more genes within a test set than the number expected from a random sampling [26], hence, we say this set is enriched with genes of the pathway, this may in turn suggest biological relevance. The statistical significance of an enrichment can be assesed by means of an hypergeometric test. In order to know if the combined regulons of our most important Trascription Master Regulators are enriched for biological pahways, an Overrepresentation Enrichment Analysis (ORA) was performed using the WebGestalt tool [27] with KEGG as the functional reference database [24]. Statistical significance threshold was set to *p* ≤ 0.05 after FDR correction.

### Pathway deregulation analysis

To determine which signal transduction pathways are the most deregulated in the breast cancer phenotype, we estimated the degree of deregulation of KEGG Signal Transduction pathways by using the *Pathifier* algorithm [28]. Pathifier assigns a score, named Pathway Deregulation Score (PDS) for each pathway in a sample from the expression status of the genes in the pathway in reference to its expression in normal tissues of the same origin. In brief, for a given pathway, a multidimensional space is defined where each dimension represents the expression level of a gene. All samples are positioned in this space according to the expression levels of all the genes in the pathway. Then, a principal curve (a smoothed curve of minimal distance to all points) is calculated and all samples are projected into it. The score corresponds to the distance of the sample projection measured over the principal curve respect of the projection of the normal tissue samples [28]. To enable comparisons between pathways a *Z*-score was calculated for each PDS and the median value for each pathway was taken [29].

### Regulon enrichment of Gene Ontology biological processes

To gain insight on how our TMRs may contribute to this phenotype, we performed an ORA for each one of their regulons against Gene Ontology (GO) [30] biological processes. Enrichments were calculated via WebGestalt [27]. Statistical significance threshold was set to *p* ≤ 0.05 after FDR correction.

## Results and discussion

From the 780 TFs in our expression matrix, 765 were involved in a total of 212,955 statistically significant interactions. MARINa detected 338 regulators in the context of breast cancer S3 File. We found that, approximately, 30 percent of the genes belong to the set that KEGG calls signal transduction pathways is regulated by GLI Family Zinc Finger 2 (GLI2), Paternally Expressed 3 (PEG3), T Box 18 (TBX18), Homeobox A5 (HOXA5), Heart And Neural Crest Derivatives Expressed 2 (HAND2), Homeobox A3 (HOXA3), Meis Homeobox 2 (MEIS2), Homeobox A2 (HOXA2), Teashirt Zinc Finger Homeobox 2 (TSHZ2) and Clock Circadian Regulator (CLOCK) from now on named ”top 10 master regulators” of signaling pathways in breast cancer (Fig. 2).

**Fig 2.**
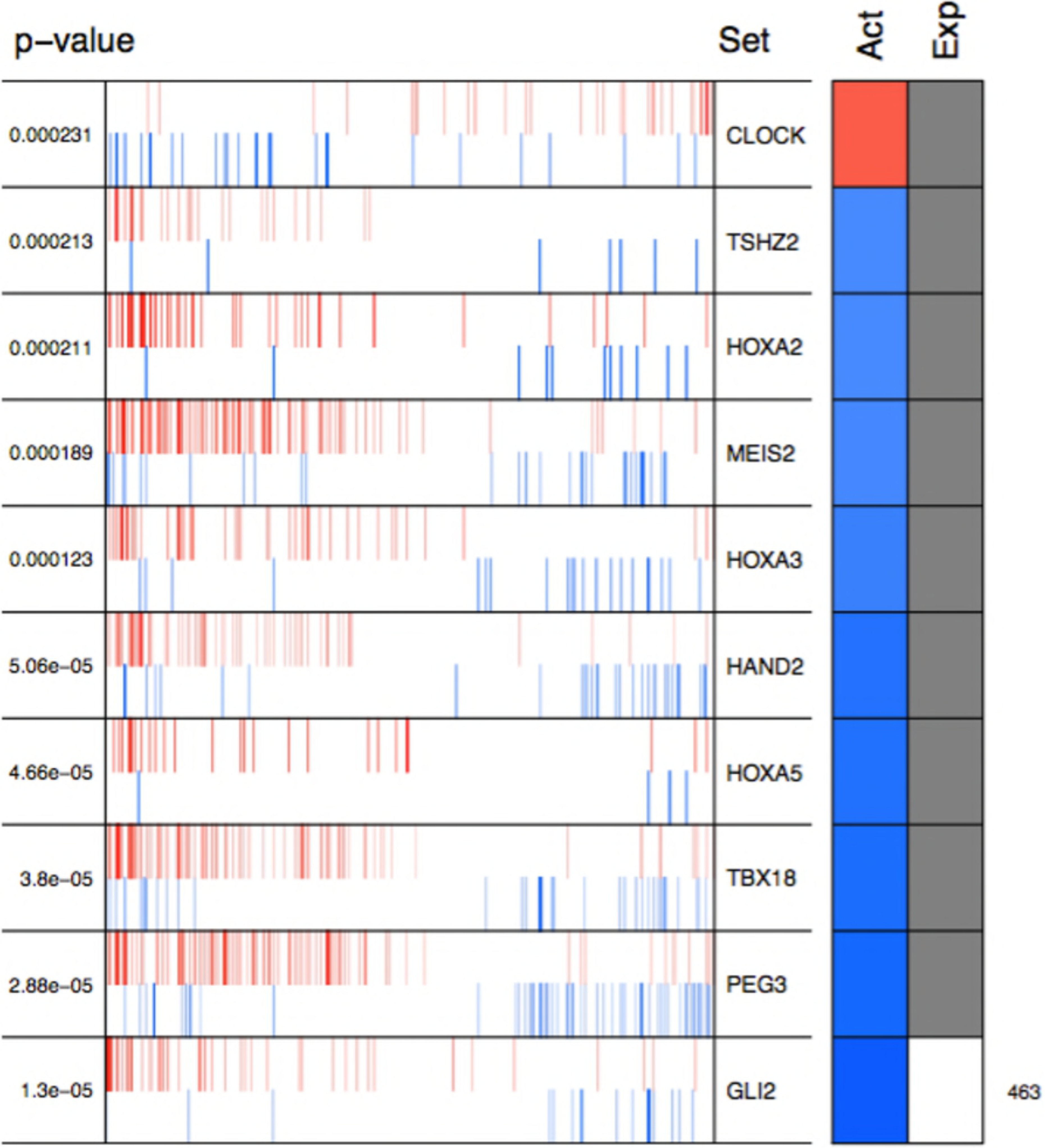
Top 10 master regulators of signal transduction pathways. These transcription factors control the genes of signal transduction pathways more differentially expressed in the tumor tissue. With the exception of CLOCK, this regulators are commonly described within the context of embryonic development, and all of them have been reported in association with cancer. The total number of genes controlled by these regulons is 412, representing almost one third of the total genes in the molecular signature. In this figure, the *p*-value is shown on the left for each of the Master regulators whose symbols are on the right. The “Act” column indicates the activity of the master regulator on its targets, the red color represents the overexpression and the blue color represents the subexpression with respect to normal tissue. “Exp” shows the expression value for each master regulator.

With the exception of CLOCK, the activity of these transcription factors over their targets is repression. However, the expression values of this regulators remain without significant change in breast cancer respect to normal mamary tissue (Fig. 2). In cancer, it has been described that some transcription factors can increase the transcription of their target genes by mechanisms independent of the increase in their gene expression [31], where various mechanisms of deregulation lead to nuclear accumulation and therefore to an increase in transcription of their target genes [32] [33] [34]. The persistent activation of certain TFs is an important event in the development of cancer [32]. These could be common mechanisms by which master regulators of signaling pathways are acting without changing their expression with respect to healthy tissue (Fig. 2).

Regulatory interactions in regulons are defined as activation if a target is overexpressed or inhibition if the target is underexpressed. The top 10 regulon-network S4 File shows a higher proportion of repression interactions over their target genes (Fig. 3). In this network GLI2 is the only TMR interacting with more than one TMR (PEG3, TBX18, HAND2, HOXA3 HOXA2 and HOXA5). All these genes together along with TSHZ2 and MEIS2 have been described as transcription factors in embryonic development. [35–43].

**Fig 3.**
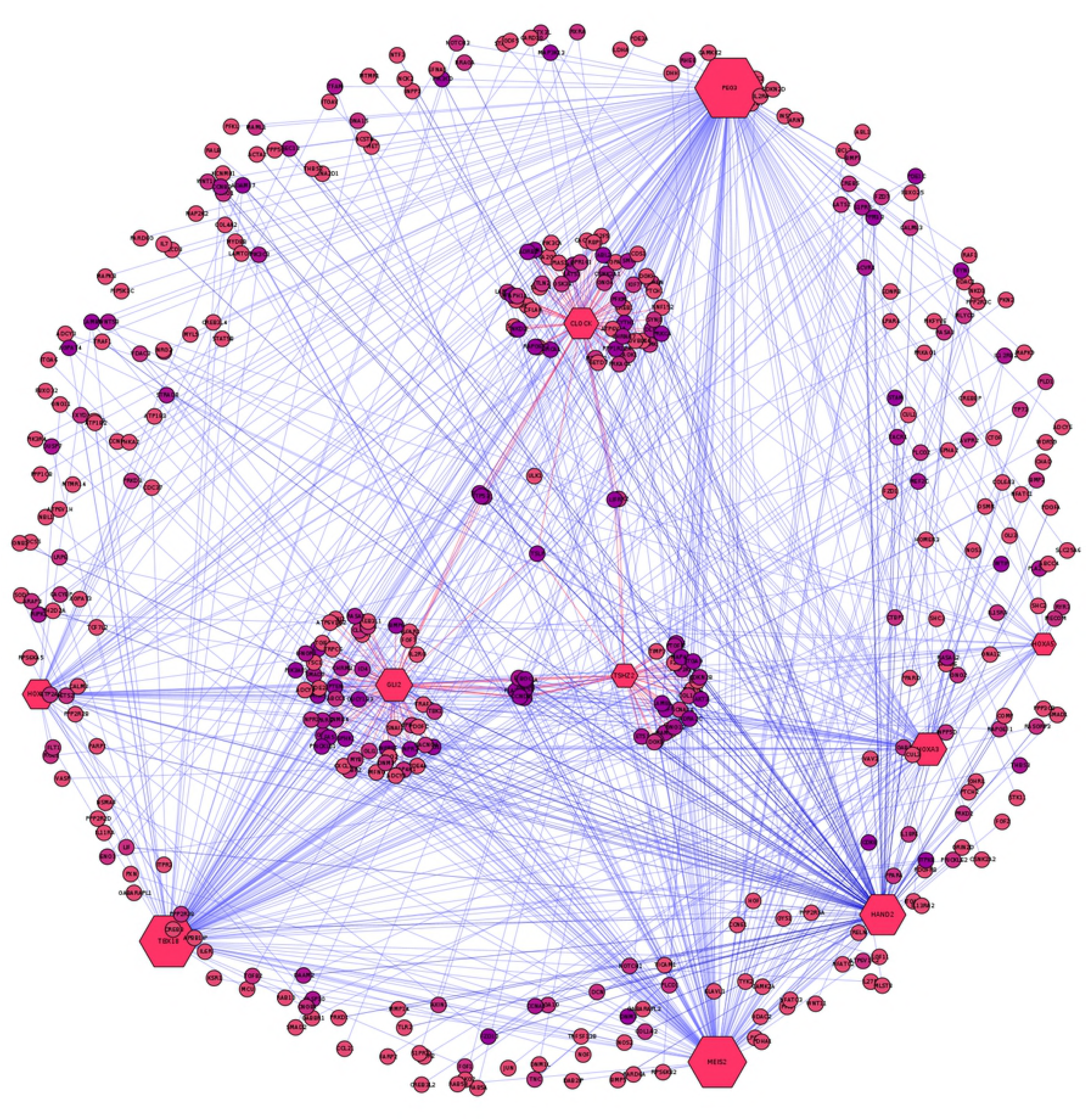
Visualization of the top 10 TMRs. Visualization of the top 10 TMRs (hexagons) and their targets (circles). TMRs show a majority of inhibition interactions of their targets (blue links). GLI2 is the TMR with the highest ES of the top 10, despite its number of interactions (hexagon size). Although it maintains activation interactions with some of its targets (red links), the majority of its interactions are inhibitory. CLOCK is the only TMR that maintains a greater proportion of activation interactions (image generated with cytoscape [44]).

### Regulon enrichment of KEGG pathways

In order to know which molecular pathways are enriched in the top 10 regulons, an enrichment analysis was made with Web Gestalt using KEGG as a reference database. The pathway with the most statistically significant enrichment was *Pathways in cancer* (hsa05200) with a coincidence of 121 genes, which reinforces the idea that the analysis does recover information from the phenotype studied.

Other pathways such as *Cell cycle* (hsa04110) and *Focal adhesion* (hsa04510) follow in the the top three enrichments. Also enriched are signaling pathways present within our molecular signature and that are known to be important in the development of cancer such as *PI3K*-*AKT signaling pathway* (hsa04151), *Phospholipase D signaling pathway* (hsa04072) and *Hedgehog signaling pathway* (hsa04340) (Table 1). These pathways seem suggestive of coordinated signalling towards survival, proliferation and differentiation.

**Table 1.** Enrichment analysis. Statistical overrepresentation analysis of KEGG pathways for the Top 10 TMR regulons network was performed with Web Gestalt. Statistical significance threshold was set to *p* ≤ 0.05 after FDR correction.

| ID | Name | # Genes | FDR |
| --- | --- | --- | --- |
| hsa05200 | Pathways in cancer | 121 | 0.000347 |
| hsa04110 | Cell cycle | 46 | 0.00226 |
| hsa04510 | Focal adhesion | 66 | 0.00386 |
| hsa05214 | Glioma | 27 | 0.00993 |
| hsa05215 | Prostate cancer | 33 | 0.0143 |
| hsa05016 | Huntington's disease | 60 | 0.0148 |
| hsa04151 | PI3K-Akt signaling pathway | 96 | 0.0148 |
| hsa04072 | Phospholipase D signaling pathway | 47 | 0.0148 |
| hsa01521 | EGFR tyrosine kinase inhibitor reistance | 30 | 0.0148 |
| hsa04340 | Hedgehog signaling pathway | 20 | 0.0148 |

### Pathway deregulation analysis

To enable comparison between different pathways by their PDS, each PDS was *Z*-score transformed and the median value of each pathway is presented in Table 2. The Hedegehog and Wnt signaling pathways showed the strongest deregulation. Meanwhile, Phosphatidylinositol and Calcium signaling pathway are the least deregulated.

**Table 2.** Pathway deregulation analysis. Median Z-scores of the PDS for the enriched KEGG Signal transduction pathways.

| Pathway | Median PDS Z-Score |
| --- | --- |
| hsa04151 PI3K-Akt signaling pathway | 0.321781930168629 |
| hsa04152 AMPK signaling pathway | 0.316562821917115 |
| hsa04340 Hedgehog signaling pathway | 0.312536134719705 |
| hsa04068 FoxO signaling pathway | 0.297611018546436 |
| hsa04015 Rap1 signaling pathway | 0.296622073957933 |
| hsa04010 MAPK signaling pathway | 0.287849136482123 |
| hsa04310 Wnt signaling pathway | 0.287489649182539 |
| hsa04014 Ras signaling pathway | 0.283094674791495 |
| hsa04330 Notch | 0.281126947323029 |
| hsa04371 Apelin signaling pathway | 0.276598032942348 |
| hsa04390 Hippo signaling pathway | 0.271001251002816 |
| hsa04350 TGF-beta signaling pathway | 0.264219294572969 |
| hsa04024 cAMP signaling pathway | 0.25893488218613 |
| hsa04668 TNF signaling pathway | 0.255446164657247 |
| hsa04012 ErbB signaling pathway | 0.25451630920054 |
| hsa04072 Phospholipase D signaling pathway | 0.250507938869976 |
| hsa04150 mTOR signaling pathway | 0.242234671909086 |
| hsa04370 VEGF signaling pathway | 0.230778782017974 |
| hsa04630 Jak-STAT signaling pathway | 0.230082497675006 |
| hsa04022 cGMP-PKG signaling pathway | 0.205220768205306 |
| hsa04064 NF-kappa B signaling pathway | 0.172261976200343 |
| hsa04066 HIF-1 signaling pathway | 0.164042143231672 |
| hsa04071 Sphingolipid signaling pathway | 0.129536138410959 |
| hsa04070 Phosphatidylinositol signaling system | 0.107851231722053 |
| hsa04020 Calcium signaling pathway | 0.0790114506475239 |

During embryonic developement signals such as morphogens and growth factors present in cell’s environment activate Signal transduction pathways that in turn induce changes within the cell [45]. In the context of cancer, pathways have been reported as permanently activated and to gain independence of the activating ligands [32].

Many of our TMRs are usually described in the context of embryonic development processes [35–43]. It is interesting to note that our TMRs and their regulons are enriched for the Hedgehog Signaling pathway. Hedgehog is an important pathway during embryonic development and in conjunction with Wnt play a role in the self-renewal of stem cells [46]. Both pathways have been previously described in cancer [3,46]. Within the TMRs that have the enriched Hedgehog pathway, it has been described that TSHZ2 forms a complex with GLI1 which functions in a coordinated manner with GLI2 and GLI3 within the Hedgehog pathway [47]. Knockout experiments of TBX18 showed a marked decrease in the Hedgehog pathway genes [48].

GLI2 regulon in the context of the top 10 regulon network. GLI2 is the only TMR that shows multiple interactions with other TMRs (six in total Figure 3). In the regulon network, genes are initially associated by means of MI but during the conversion to regulons directionality is assigned from TF to other genes. Whenever an interaction between two TFs is present, directionality is not resolved. GLI2, together with GLI1, GLI3 [49] and TSHZ2 (another of our TMRs) [47] are important effector molecules activated within the Hedgehog pathway that modulate dedifferentiation and differentiation processes during embryonic development [42,50]. Therefore, this TMR may be interesting in the context of the master effector of the Hedgehog pathway which is one of the most represented here.

Another interesting result arises from the observation that the PI3K-AKT and Hedgehog signaling pathways have been reported in association with stemness and cell differentiation processes. Both pathways play a role during embryonic development and in the maintenance of adult tissues. Hedgehog plays a role in epithelium maintenance, and is necessary to regulate the presence and number of stem cells [3], while activation of the PI3K-AKT pathway promotes survival growth and proliferation [51].

### Enrichment of each regulon in GO processes

The most significantly enriched processes of each TMR regulon are presented in Table 3. It is interesting that, for enriched GO biological processes obtained from the molecular signature of the signal tranduction pathways, the top places are occupied by embryonic development related processes. These results are in line with the hypothesis of tumors are described as aberrations of growth, differentiation, and organization of cell populations. These are basic processes that are tightly coordinated and controlled during embryogenesis as well as in adult tissues [6]. The oncogerminative theory of cancer development (OTCD) [6] suggests that cancer arises due to aberrant expression of developmental genes. According to this theory, tumor formation is a dynamic self-organizing process that mimics the process of early embryo development. The malignant transformation of somatic cells, which is a result of gene mutations combined with epigenetic dysregulation, ultimately results in somatic cells being reprogrammed into immortal cells that mimic germline cells. These mimics are termed “cancer stem cells” or “oncogerminative cells” [6,52].

**Table 3.** First significant enrichments of Gene Ontology biological processes per regulon. The first ten regulons enrich more biological processes related to embryonic development (ten out of fifteen, in purple), in blue are the processes related to cell cycle and proliferation (three enriched processes) and in orange those referring to organization of the extracellular matrix (two).

| Regulon | Enriched GO processes | ID | FDR |
| --- | --- | --- | --- |
| CLOCK | Mitotic cell cycle | GO:0000278 | 1.39E-02 |
| GLI2 | Regulation of cell differentiation | GO:0045595 | 1.22E-05 |
| HAND2 | Cardiovascular system development | GO:0072358 | 4.31E-06 |
| HAND2 | Vasculature development | GO:0001944 | 4.31E-06 |
| HOXA3 | Tube development | GO:0035295 | 8.94E-05 |
| HOXA5 | Proximal/distal pattern formation | GO:0009954 | 1.69E-02 |
| HOXA5 | Anterior/posterior pattern specification | GO:0009952 | 1.69E-02 |
| HOXA5 | Skeletal system development | GO:0001501 | 1.69E-02 |
| MEIS2 | Animal organ morphogenesis | GO:0009887 | 5.29E-08 |
| PEG3 | Cell cycle | GO:0007049 | 8.72E-08 |
| TBX18 | Tissue development | GO:0009888 | 3.59E-05 |
| TBX18 | Blood vessel development | GO:0001568 | 3.59E-05 |
| TSHZ2 | Regulation of cell proliferation | GO:0042127 | 4.91E-02 |
| TSHZ2 | Extracellular matrix organization | GO:0030198 | 4.91E-02 |
| TSHZ2 | Extracellular structure organization | GO:0043062 | 4.91E-02 |

In humans Homeobox A family cluster consists of eleven genes (HOXA1, HOXA2, HOXA3, HOXA4, HOXA5, HOXA6, HOXA7, HOXA9, HOXA10, HOXA11, HOXA13). Although HOXA genes code for proteins with transcription factor activity, these are not typically considered as components of signal transduction pathways. HOXA TFs act not only as transcriptional activators in cancers but also as transcriptional repressors [53], thus, both upregulation and downregulation of the members of this family may be critical for promotion of carcinogenesis. Many HOXA genes (HOXA1, A2, A3, A5 and A9) have been shown to have significantly lower expression levels in cancerous tissues compared to non-cancerous tissues. In human breast cancer cells, HOXA5 was observed to activate the p53 tumor suppressor gene promoter [54]. Expression of HOXA5 in breast cancer cells expressing wild-type p53 led to apoptosis while those lacking the p53 gene did not [54,55]. Furthermore, the HOXA5 promoter region was methylated in 80 % of p53-negative breast cancer specimens. [54]. This aberrant regulation of HOX genes in cancer indicates that HOX transcriptional mechanisms are integral to a network of regulatory mechanisms involved in normal adult tissue homeostasis. [52]. Our results show that HOXA members are included in all of our 10 TMR regulons Table 4.

**Table 4.** HOXA Family Genes present in Top TMR regulons. Numerous HOXA family members are part of the top TMR regulons and significant *p* value was found in all cases. Hypergeometric test parameters are: population size **N** = 15802 genes in the expression matrix, number of successes in population **M** = 10 eleven human HOXA genes in expression matrix, sample size **s** is the regulon size and number of successes **k** is the number of HOXA genes present in the regulon. HOXA13 was not present in any of the top regulons.

| HOXA | CLOCK | GLI2 | HAND2 | HOXA2 | HOXA3 | HOXA5 | MEIS2 | PEG3 | TBX18 | TSHZ2 |
| --- | --- | --- | --- | --- | --- | --- | --- | --- | --- | --- |
| HOXA1 | 0 | 1 | 1 | 1 | 1 | 1 | 1 | 1 | 1 | 0 |
| HOXA2 | 1 | 1 | 0 | 1 | 1 | 1 | 1 | 1 | 0 | 0 |
| HOXA3 | 1 | 1 | 1 | 1 | 1 | 1 | 1 | 1 | 1 | 1 |
| HOXA4 | 1 | 1 | 1 | 1 | 1 | 1 | 1 | 1 | 1 | 1 |
| HOXA5 | 0 | 1 | 0 | 1 | 1 | 0 | 1 | 1 | 1 | 0 |
| HOXA6 | 0 | 1 | 0 | 0 | 0 | 0 | 0 | 1 | 0 | 0 |
| HOXA7 | 1 | 1 | 1 | 1 | 1 | 1 | 1 | 1 | 1 | 0 |
| HOXA9 | 0 | 1 | 0 | 1 | 1 | 1 | 1 | 0 | 1 | 1 |
| HOXA10 | 1 | 1 | 0 | 1 | 1 | 1 | 1 | 1 | 1 | 1 |
| HOXA11 | 0 | 0 | 1 | 1 | 1 | 1 | 0 | 1 | 1 | 0 |
| Total | 5 | 9 | 5 | 9 | 9 | 8 | 8 | 9 | 8 | 4 |
| $p$ value | 8.19e-5 | 9.78e-13 | 6.48e-5 | 8.0e-13 | 1.64e-11 | 4.72e-13 | 1.22e-7 | 1.69e-7 | 5.13e-8 | 5.25e-5 |

## Conclusion

Through the generation of a signal transduction-focused molecular signature we identified the top 10 TMRs that, in combination regulate up to 30% of the molecular signature genes. A further analysis of the gene sets conformed by the top TMRs and associated regulons pointed out to the PI3K-AKT pathway, which is associated to cell survival and proliferation, and to the AMPk pathway which is involved in the cellular energetic balance as the most deregulated.

Nine out of our ten TMRs are recognized for taking part in embryonic development associated processes [35–43]. In consonance with this, the Hedgehog signaling pathway was shown as enriched and highly deregulated. Further individual GO biological function enrichments for each TMR associated regulons showed six out of ten significantly enriched in embryonic development related processes. Given the functional and gene composition overlap between the regulons, it appears as an indication of the presence of a gene regulation module where signal transduction pathways are cooperatively regulated by a set of TMRs in a way that embryonic development processes are subverted in favor of tumor development.

Signal transduction pathways are characterized by taking external signals to generate intracellular changes. The cellular functions enriched in the regulons controlled by the top TMRs associated to these pathways are focused around embryonic development processes. Because of this, we suggest that the signaling pathways could be deregulated through genetic mechanisms such as mutations, or that are receiving signals from the external environment that lead to aberrant activation of signaling pathways typical of embryonic development to give the breast cnacer cell its distinctive proliferative, survival and angiogenesis capabilities.

Hence, by analyzing the activity of transcriptional master regulators over pathway-prioritized genesets, it is possible to look at process-specific regulatory patterns that help us to uncover specific biological functions. This in turn may open up novel ways of inquiry, useful to develop system-wide semi-mechanistic descriptions of complex phenotypes, such as cancer.

## Supporting information

**S1 File. Transcription factor list.** List of transcription factors taken from the TFcheckpoint database.

**S2 File. List of genes in the molecular signature.** List of genes in our expression matrix that are in KEGG Signal Transduction category pathways.

**S3 File. Inferred TMRs.** Table of inferred TMRs with regulon size, nes, p-value and FDR-corrected p-value.

**S4 File. Top 10 TMR regulons network.** Network file in .sif (node-interaction-node list) format of the top 10 TMR regulons.

## Acknowledgments

This paper constitutes a partial fulfilment of the Graduate Program in Biological Sciences of the National Autonomous University of México (UNAM). D. Tapia-Carrillo acknowledges the scholarship and financial support provided by the National Council of Science and Technology (CONACyT), and UNAM. This work was supported by CONACYT (grant no. 285544/2016), as well as by federal funding from the National Institute of Genomic Medicine (Mexico). Additional support has been granted by the Laboratorio Nacional de Ciencias de la Complejidad (grant no. 232647/2014 CONACYT). EH-L is recipient of the 2016 Marcos Moshinsky Fellowship in the Physical Sciences.

